# Improving allele-specific epigenomic signal coverage by *10-fold* using Hidden Markov Modeling and Machine Learning

**DOI:** 10.1101/2024.05.23.595536

**Authors:** Emmanuel LP Dumont, Ali Janati, Moumita Bhattacharya, Jean-Baptiste Jeannin, Catherine Do

**Affiliations:** Hackensack Meridian Center for Discovery and Innovation; Columbia University; Netflix Research; University of Michigan; New York University Langone Health

## Abstract

Allele-specific epigenomic signals refer to differences in epigenomic patterns between the two copies, or “alleles,” of a DNA region inherited from each parent. Epigenomic patterns are defined as alterations of the DNA sequence (e.g., chemical) without modifying the underlying DNA sequence (which would be referred to as “mutations”). Mapping allele-specific epigenomic signals across a genome is crucial, as some can influence gene expression, disease susceptibility, and developmental processes. However, identifying allele-specific epigenomic patterns across an entire genome is limited by the average read length (50-150 nucleotides) of short-read sequencing technologies, which are the most widely-used and affordable whole genome sequencing methods, and by the 99.9% similarity in the DNA sequences inherited from each parent. These limitations restrict the assessment of allele-specific signals to approximately 10% of the genome, potentially overlooking critical regulatory regions. In this paper, we present a highly effective machine-learning approach based on variational hidden Markov modeling, which enables the detection of allele-specific epigenomic signals across the entire genome, resulting in a 10-fold improvement in genomic coverage compared to state-of-the-art methods. We demonstrate our method on DNA methylation, a critical epigenomic regulatory signal.

## 1 Introduction

### The role of allele-specific epigenomic signals in the regulation of the genome

In individuals, the Deoxyribonucleic acid sequence (**DNA**) comprises three billion nucleotides. These nucleotides can differ by their nitrogenous bases, which are adenine (A), cytosine (C), guanine (G), or thymine (T). These four bases’ specific order or sequence along the DNA molecule encodes the genetic instructions. The DNA is divided into 23 pairs of chromosomes. The two chromosomes in each pair are inherited from the individual’s parents and are approximately 99.9% similar in their sequence. Each parental copy of the DNA is called an “allele” (paternal or maternal). The primary role of DNA is to be transcribed into RNA, which is then translated into proteins. However, only approximately 1% of the genome codes for proteins (these regions are called “genes” and are said to be “coding”). In the remaining 99% of the genome (also called “non-coding”), a partially unidentified set of regions has been found to regulate gene expression. Discovering and understanding how these regions indirectly impact protein production is an active area of scientific research at the intersection of molecular biology and computational genomics, falling under the umbrella of “epigenetics.” Epigenetics refers to additional layers of regulation that can influence gene expression without altering or mutating the underlying DNA sequence. These layers of regulation play an essential role in cell differentiation, disease development, etc. [1] There are three known epigenomic signals: DNA methylation, histone modifications, and chromatin accessibility, shown in Figure 1. To flag the genome with these epigenomic markers, biologists use specific molecular biology assays to prepare the DNA. Finally, one distinctive and important feature of allele-specific epigenomic signals is that they occur in only 1% of the genomic regions that can be assessed for this imbalance [2]. This strong imbalance in classes will pose specific challenges in predicting them using machine learning.

**Figure 1:**
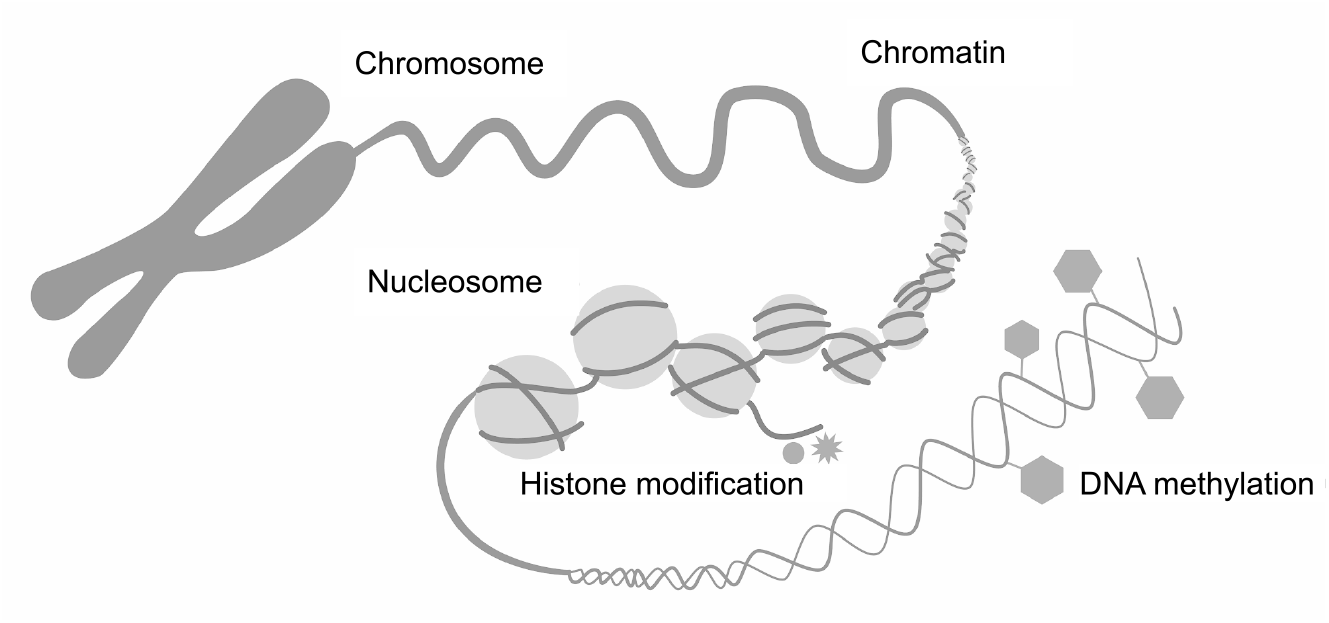
Epigenetics refers to mechanisms that regulate DNA expression without altering the DNA sequence. Histone modifications serve to expand or contract the chromatin into “open” or “closed” configurations. DNA methylation refers to the addition of methyl groups to cytosines positioned within cytosine-phosphate-guanine (**CpG**) sequences across the genome. Figure adapted from https://www.zymoresearch.com/

### Whole genome sequencing

After the DNA is extracted from a sample (e.g., blood), a “library” of randomly fragmented DNA is generated and then sequenced. Those sequences are between 50 -150 nucleotides long, depending on the biology protocol. Multiple sequences, or “reads”, are generated for a given genomic region to correct for sequencing errors later on. The average number of reads per nucleotide position in the sequenced genome is called “coverage” and ranges from 10 to about 100. These reads are then “aligned” to the reference genome, enabling researchers to map them and find their coordinates along the DNA sequence. When assessing allele-specific events, higher coverage is critical since the sequencing reads are separated between the two copies of the chromosomes inherited from the individual’s parents and analyzed separately. However, because these copies are 99.9% identical in their sequence, 90% of the sequencing reads cannot be separated by their chromosome of origin, preventing researchers from assessing allele-specific epigenomic signals in most of the genome. This severely hinders our understanding of epigenomic regulations. To overcome this, long-read sequencing is an active research area but suffers from other limitations and is still not widely used today because of the higher cost [3].

### Contributions

Our focus is on applying time-series machine-learning techniques to epigenomic data to decipher allele-specific epigenomic signal across a whole genome. The main contributions of the paper can be summarized as follows:

- We present an innovative *directional fractional content* variable derived from the sequencing reads and derive meaningful features from it using a Gaussian variational Hidden Markov Model (**HMM**).
- Our approach circumvents the current technological limitations of short-read sequencing technologies, which are the most widely used today. It enables researchers to assess allele-specific epigenomic signals across the whole genome, as opposed to approximately 10% of the genome for current state-of-the-art methods, representing a 10-fold increase in genome coverage.
- When applied to methylation, we show that our best model’s predictions generalize to an F1 score (harmonic mean of precision and recall, ranging from 0 to 1) of 0.995 for the absence of allele-specific methylation and 0.712 for the presence of allele-specific methylation. Our best model’s macro average F1 score (arithmetic mean of the classes’ F1 scores) is 0.854.
- Traditional methods for identifying allele-specific signals are complex and computationally expensive because they require several additional bioinformatics steps (e.g., “variant calling” to separate sequencing reads based on their chromosome of origin).

## 2 Methodology

### 2.1 Current state-of-the-art methods

Identifying allele-specific epigenomic signals is an active field of research in computational genomics [4, 5, 6, 7, 8]. Because there is no clear definition of an imbalance of an epigenomic signal between two alleles, several methods exist to assess allele-specific epigenomic signals. These methods compete in identifying “biologically true” imbalances, which are often later validated using biological cross-validations (e.g., CRISPR-Cas9 genetic editing to reproduce the imbalance). Despite the lack of a unified definition of allele-specific epigenomic signal, all these methods, by definition, require at least a single nucleotide change between the two alleles so that the sequencing reads can be assigned to a specific allele. These single nucleotide changes are called single nucleotide polymorphisms (**SNP**). Because of this design requirement, these methods only cover approximately 10% of the genome because most sequencing reads are not long enough to overlap an SNP (they are 50-150 nucleotides long on average), and SNPs are relatively rare (two parental chromosomes are 99.9% similar in their sequence). Finally, allele-specific epigenomic signals are relatively rare, as they were found to occur in about 1% of the regions that can be assessed [2]. This severe class imbalance poses additional challenges in predicting it accurately.

To address this severe limitation in genome coverage of allele-specific epigenomic signal analysis, researchers tried identifying allele-specific signals using the signal pattern among the sequencing reads. Building on Fang et al. [9] and Peng and Ecker [10], Wu et al. [11] proposed a nonparametric Bayesian clustering method applied to the epigenomic signal of methylation. Their method group sequencing reads into 2 clusters based on their methylation content, using a fast Dirichlet process-mixture search method, and compute if the average methylation across both clusters is statistically significant. These methods, however, do not offer a simple baseline approach using simple concepts and statistics to compare themselves to, which we present in this paper (See our methods), casting doubt on the added efficacy of their method. Also, these methods rely on a simple approach of assessing allele-specific imbalance without considering the direction of imbalance for each CpG, leading to many false positives in their predictions.

Not directly related to our work but still relevant, Angermueller et al. [12] have used convolutional neural networks to predict the methylation status of a single CpG in single-cell analysis. Their method does not suffer from the class imbalance we are facing, because about 60% of CpGs are methylated, and cannot identify allele-specific patterns. He et al. [13] used machine learning to predict allele-specific methylation, but one of their features is the presence of an SNP, making it unclear why one would use machine learning if they had access to an SNP, enabling them to use any state-of-the-art method utilizing the presence of an SNP among the reads.

### 2.2 Overview of our method

We develop a framework to identify allele-specific epigenomic signals across entire genomes. Because each epigenomic signal requires a complex, highly specific and computationally intensive bioinformatics pipeline to align the sequencing reads flagged with their epigenomic signal, we present the results of our methodology for methylation only. However, our method can be used for all epigenomic signals provided that sequencing reads are flagged with a fractional content of the epigenomic signal (e.g., Fiber sequencing for histone modification) and not a binary outcome (presence or not of the epigenetic signal in the read). Our framework takes advantage of the sequential nature of the DNA sequence, variational HMMs, and publicly available epigenomic-marked genomes. HMMs were introduced by Baum and Petrie [14] for speech processing, and the variational Bayesian implementation of HMM was first introduced by MacKay [15]. The data we used for training, validation, and testing is publicly available on the ENCODE website, a public repository of whole genomes marked with epigenomic signals [16]. Our model infrastructure builds on a simple idea: to sort sequencing reads by the fractional content of their epigenomic signals to infer the directional fractional content (**DFC**) of the signal, which is a number between -1 and 1 but with a distribution skewed toward the interval [0, 1] driven by the read sorting, as shown in Figure 3. This DFC sequence is then trained on a Gaussian Variational HMM with three hidden states, which we interpret as positive imbalance, negative imbalance, and no imbalance on each signal along the DNA sequence. After computing each signal’s most likely hidden state along the DNA sequence, we computed several features for each genomic region (number of states, transition probability between each state, entropy, *etc*.), which we used in a tree-based classification algorithm.

We also present a simple baseline method against which to compare our work (See Results section). Our baseline algorithm sorts the sequencing reads by their content in the epigenetic signal and applies the same criteria shown in Figure 2, leading to a binary prediction of allele-specific signals. In DNA sequencing, unless a biology protocol introduces a known bias on purpose, it is generally assumed that the sequencing reads sampled across any given genomic region are equally likely to come from either of the two maternal and parental chromosomes. This assumption is central to many analytical methods in genomics, including variant calling.

**Figure 2:**
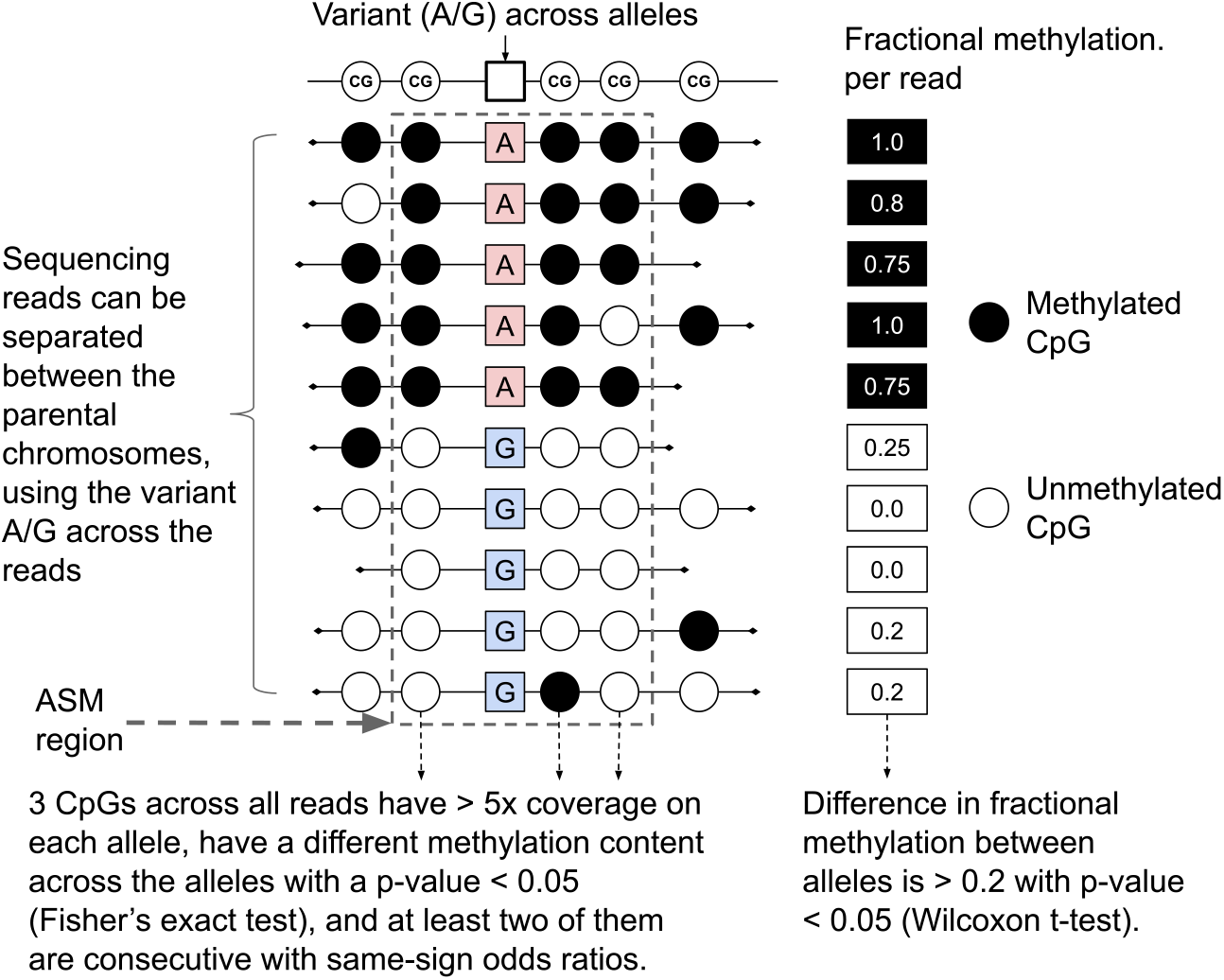
Each research group has developed its method of detecting allele-specific signals. Here, we show the method developed by Do et al. [2] to assess allele-specific methylation, which we used as a label in our modeling. Only the sequences Citosine-phosphate-Guanine (**CpG**) are shown because only these sequences can be methylated in DNA. Figure adapted from Dumont et al. [6]

Finally, we have also implemented several sequential models (transformers and recurrent neural networks) to the DFC sequence, which we will show do not perform as well as training a tree-based algorithm on features derived from the hidden states of a 3-state Gaussian Variational Hidden Markov model.

### 2.3 Derivation of the Directional Fractional Content (DFC)

For a given genomic region with *N* sequencing reads overlapping it (*N* is also the coverage for that region), we sort the reads by their content of fractional epigenomic mark (from *k* = 1 to *k* = *N* ), and we define the DFC at the *i*th position as follows, where *M* is the epigenomic signal:

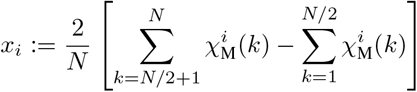

Where 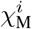 is the indicator function of the epigenetic signal *M* at the position *i*. In Figure 3, we illustrate our methodology and present the outcome of the trained HMM for DNA methylation.

**Figure 3:**
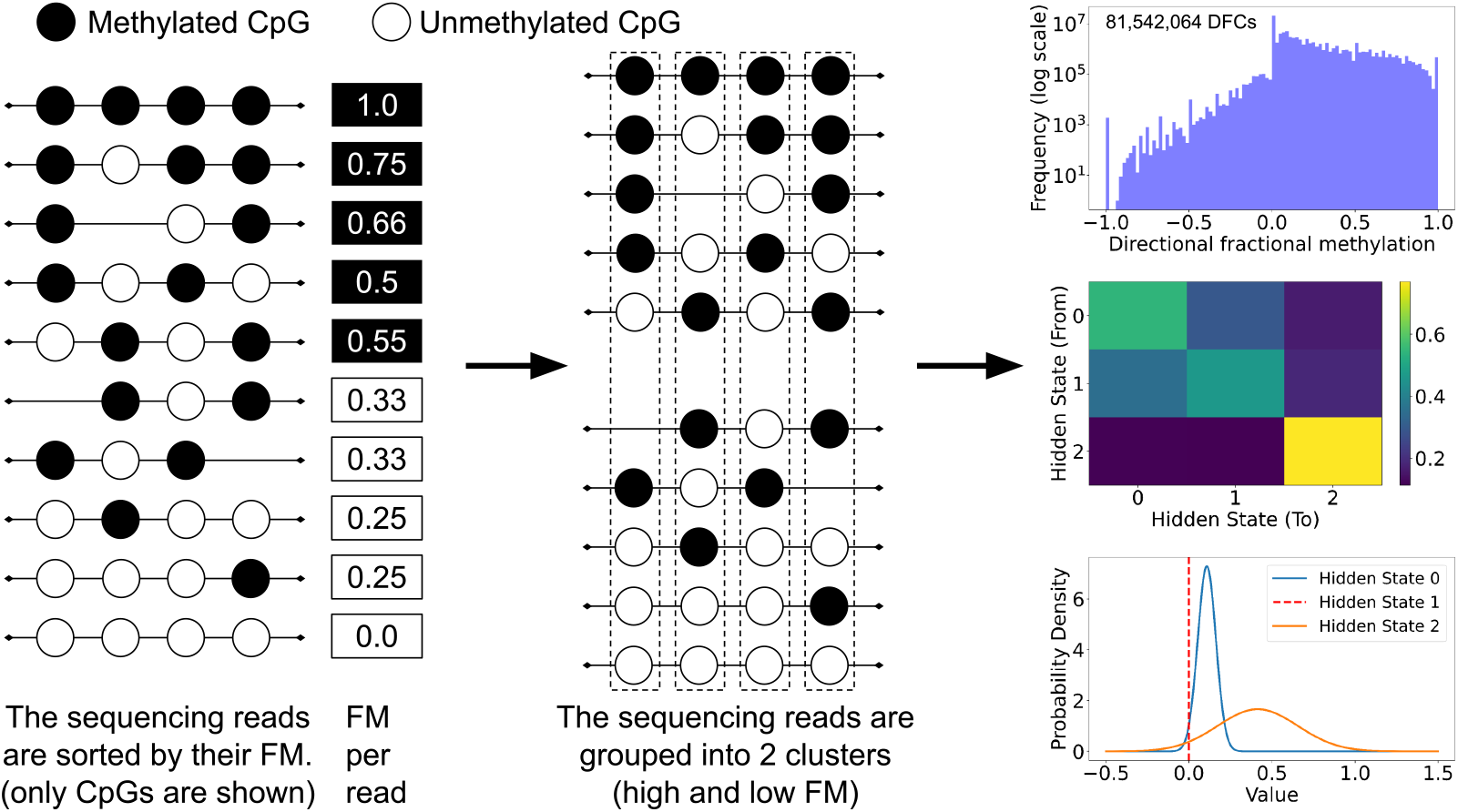
Illustration of the derivation for the DFC for methylation (which we call directional fractional methylation or directional FM). We first sort sequencing reads by their FM (left panel). We only represent CpGs in the sequence since only CpGs can be methylated. Note that some reads may not overlap all CpGs in the region. We then compute each CpG’s directional FM (middle panel) by splitting the reads into 2 groups (high and low FM) and then fit a Gaussian variational HMM with three independent hidden states to all concatenated DFCs. We picked three hidden states to have them represent a positive imbalance, negative imbalance, and no imbalance in CpG methylation across the two groups of sequencing reads. Finally, we used 11 million DFCs to fit the Variational Gaussian HMM after randomly picking a subset of genomic regions - there are 81.5 million DFCs in the training set in total.

### 2.4 Gaussian Variational HMM

We used a Hidden Markov Model with three independent hidden states {*S*_1_, *S*_2_, *S*_3_} and not a mixture of hidden states. The reason for this choice is anchored in biology. Each hidden state is intended to represent the individual state of individual signals along the DNA sequence: positively imbalanced, negatively imbalanced, and no imbalance. As a result, the signal is not expected to be built as a mixture of hidden states. Each state *S*_*j*_ is modeled by a Gaussian distribution in a one-dimensional space, with mean *µ*_*j*_ and variance 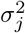. The probability of observing *x*_*i*_ given the state *S*_*j*_ is given by the following formula:

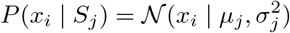

where the one-dimensional Gaussian distribution is given by:

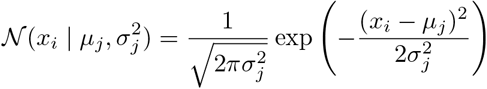

Variational Inference provides an alternative approach to Expectation-Maximization (**EM**) that turns the estimation of the posterior distributions of latent variables into an optimization problem. It involves approximating the true posterior with a simpler distribution by minimizing the Kullback-Leibler divergence between the true posterior and the approximate distribution. Variational inference can be more robust than EM in scenarios where the EM algorithm might get stuck in local optima.

Given a sequence of observations of DFCs **x** = (*x*_1_, …, *x*_*n*_) and a corresponding sequence of independent hidden states **z** = (*z*_1_, …, *z*_*n*_) where *z*_*i*_ ∈ {*S*_1_, *S*_2_, *S*_3_}, the joint probability of the observations and the state sequence can be expressed as follows:

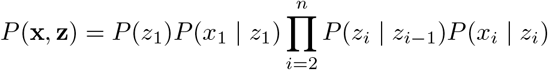

Where: *P* (*z*_1_) is the initial probability of starting in state *S*_1_, *P* (*x*_1_ | *z*_1_) is the probability of observing *x*_1_ given the initial state *z*_1_, *P* (*z*_*i*_ | *z*_*i−*1_) is the transition probability from state *z*_*i−*1_ to state *z*_*i*_, and *P* (*x*_*i*_ | *z*_*i*_) is the probability of observing *x*_*i*_ given state *z*_*i*_.

### 2.5 Supervised Learning

For variational Gaussian HMMs, we applied the following steps:

- We fitted the Variational Gaussian HMM on a random subset of the genomic regions from the training dataset. We used over 11 million DFCs to train our HMM (among 81 million DFCs available).
- For all datasets (training, validation, test), we inferred the hidden state for each DFC in each genomic region, derived features for each genomic region based on these states (number and proportion of each hidden state, number of transitions and transition probability from one hidden state to another, mean and variance of the duration of each hidden state, and entropy state distribution, first and last hidden state). We also applied kernel functions to the distribution of CpG distances, coverage, fractional methylation, and sequencing reads fractional methylation.
- We trained each algorithm in each algorithm class (random forest, gradient-boosted tree combined with a type of Gaussian HMM) using the samples in the training set and across a grid of hyper-parameters using a Random Search.
- We picked the best model in each class by using the sum of the F1 scores of each class on the validation set.
- We retrained the best model across the training and validation sets.
- We evaluated the performance of each class of models on the test dataset.

For transformer and recurrent neural networks (**RNN**) architectures, we applied the following steps:

- We trained several architectures across a grid of hyper-parameters using a Random Search.
- We picked the best model in each class by using the sum of the F1 scores of each class on the validation set.
- We retrained the best model across the training and validation sets.
- We evaluated the performance of each class of models on the test dataset.

In our supervised algorithms, we used the Binary Cross-Entropy (**BCE**) with logits for our loss function. This loss function combines a sigmoid activation with the BCE loss in a single, numerically stable, and efficient formulation. Traditional cross-entropy loss typically requires the outputs to be passed through a sigmoid function to convert logits to probabilities before computing the loss. This additional step can lead to numerical instability, especially when logits are very large or small, which is our case given the highly imbalanced dataset. During training, we use batch sizes of 1,024 genomic regions to ensure that each batch includes several examples of the minority class. By integrating multiple instances of these minority class examples in every batch, we facilitate a more consistent and effective learning process across all classes.

## 3 Dataset

### 3.1 Overview

For this work, we used the ENCODE repository, a publicly available database of cell lines and human tissues analyzed across several epigenetic marks, including methylation [16]. We used six whole genomes for training, three for validation, and three for testing. We assessed 715 million sequencing reads spanning 36 billion nucleotides across all samples. This dataset has about 9 billion rows and is 2 TB in size across all samples. Due to the extensive size of the dataset, we developed a specialized cloud-based infrastructure for data processing. For more details, see our GitHub repository.

### 3.2 Data preparation and labeling

We used the GRCh37/hg19 reference genome, which we split into genomic regions of 250 nucleotides long, and we kept only the regions with at least 2 CpGs that do not overlap with ENCODE’s “blacklist” regions (regions known to have anomalous sequencing data) [17]. For each sample, we performed an alignment to the reference genome and extracted the methylation status of each CpG and its corresponding read ID. We then grouped these rows per sample per genomic region. We only retained genomic regions that had at least three CpGs, as required by our labeling method [6]. To create the DFC for each CpG, we requested to have at least 20 reads per genomic region and requested that each retained sequencing read included at least 50% of the total number of CpGs found in each genomic region.

In total, across all samples, we have 36,710,628 genomic regions. Using Dumont et al. [6], as illustrated in Figure 2, we were able to evaluate 2,926,215 regions (7.9%) for allele-specific methylation (**ASM**), where there was an overlapping SNP to differentiate the sequencing reads between the parental chromosomes. Among these, there are 35,416 regions (1.2%) where we found ASM. We were able to build our dataset of features in 29,806,134 genomic regions, representing a 10-fold improvement over the 2.9 million regions where we could flag regions with the presence of ASM, using the state-of-the-art method we had previously developed [6].

## 4 Results

We first established a baseline prediction in genomic regions. To do that, we sorted the reads by their fractional content in methylation and applied the same statistical tests and criteria described in Figure 2. In Table 1, where we present our results, this method is referred to as “not ML”, ML referring to Machine Learning. In the test dataset, which we used only to assess the generalization error of each model class, we have 528,823 regions without ASM and 7,966 regions with ASM, for a total of 536,789 genomic regions. We also implemented other Variational Gaussian HMMs (e.g., 5 hidden states with a tied covariate matrix) and a variational autoencoder to build a latent vector of the sequence of DFCs. These results are not shown in Table 1. Early experiments on variational autoencoders did not show promising results, however.

**Table 1:** Model results. VGHMM stands for Variational Gaussian Hidden Markov Model, RNN stands for recurrent neural network, and XGBoost stands for Gradient-Boosted tree. All VGHMM were fitted using 3 independent hidden states. A supervised approach using a random forest classifier on features derived from the three hidden states of the VGHMM provides the best result.

| **Model Description** |  |  |  |  |  |
| --- | --- | --- | --- | --- | --- |
| Model category | Not ML | **Random Forest** | XGBoost | Transformer | RNN |
| Underlying structure | Statistical tests | **VGHMM** | VGHMM | N/A | N/A |
| **Confusion matrix** |  |  |  |  |  |
| True Positive | 7,768 | **6,250** | 7,299 | 0 | 6,486 |
| True Negatives | 427,691 | **525,489** | 522,587 | 528,370 | 488,589 |
| False Positives | 100,678 | **2,880** | 5,782 | 0 | 39,781 |
| False Negatives | 652 | **2,170** | 1,121 | 8,420 | 1,934 |
| **Classification report** |  |  |  |  |  |
| *Precision* |  |  |  |  |  |
| Class 0 | 0.998 | **0.996** | 0.998 | 0.984 | 0.996 |
| Class 1 | 0.072 | **0.684** | 0.558 | 0.0 | 0.140 |
| Macro average | 0.535 | **0.840** | 0.778 | 0.492 | 0.568 |
| *Recall* |  |  |  |  |  |
| Class 0 | 0.809 | **0.995** | 0.989 | 1.0 | 0.925 |
| Class 1 | 0.923 | **0.743** | 0.867 | 0.0 | 0.770 |
| Macro average | 0.866 | **0.868** | 0.928 | 0.5 | 0.848 |
| *F1 score* |  |  |  |  |  |
| Class 0 | 0.894 | **0.995** | 0.993 | 0.992 | 0.959 |
| Class 1 | 0.133 | **0.712** | 0.679 | 0.0 | 0.237 |
| Macro average | 0.514 | **0.854** | 0.836 | 0.496 | 0.598 |
| *Sum of F1s across classes* | 1.027 | **1.707** | 1.672 | 0.992 | 1.196 |

Interestingly, despite employing Random Search to optimize their configurations, including the use of high-dimensional feature vectors through deep networks, RNNs and transformers showed negligible improvement or even underperformed compared to the non-machine learning approach in our tests. We believe that the simple representation of three independent hidden states reflects well the biological nature of DFCs in the context of allele-specific analysis (Positive Imbalance: More methylation on one allele compared to the other; Negative Imbalance: Less methylation on one allele than the other; No Imbalance: Equal methylation across alleles). This intuitive approach turned out to yield the best results in our work.

## 5 Conclusion, Limitations, and Future Work

We present DeepASM, a model architecture that leverages the time-series nature of genomic data and variational HMM to assess allele-specific epigenomic signals in entire genomes rather than being limited to genomic regions where an SNP is present. Our method represents a significant improvement over non-machine learning methods with a generalization macro average F1 score of 1.707. Our infrastructure was built for methylation but can be used for other epigenomic signals, provided that sequencing reads are flagged with a fractional content of the epigenomic signal. This represents a 10-fold improvement over traditional methods in genome coverage of allele-specific epigenomic signals.

However, our approach has limitations. The primary limitation of our method is that it tends to predict imbalances as predominantly positive due to the way sequencing reads are sorted. As a result, two different genomic regions where our predictions show an allele-specific epigenomic signal could be in opposite directions, which our method would not predict. Another technical limitation is that we requested that sequencing reads carry at least 50% of the signal found in the genomic region. This resulted in a loss of 19% of the genomic regions where allele-specific epigenomic signals cannot be predicted. Future work can focus on being less stringent and building a better model to address these regions. Another limitation is the lack of a computer-vision approach to the pattern of epigenomic signals across all reads in a genomic region. This approach is expected to be much more computationally expensive but should be assessed. Finally, the variational Gaussian HMM should be fitted on all 81 million DFCs in the training set rather than selecting random genomic regions amounting to 11 million DFCs.

## Acknowledgments and Disclosure of Funding

We are grateful to the Hackensack Meridian Center for Discovery and Innovation and the BD Foundation for their generous funding and to Benjamin Tycko for his insightful comments on our manuscript.

